# Mechanism of ligand recognition by human ACE2 receptor

**DOI:** 10.1101/2020.10.30.362749

**Authors:** Apurba Bhattarai, Shristi Pawnikar, Yinglong Miao

## Abstract

Angiotensin converting enzyme 2 (ACE2) plays a key role in renin-angiotensin system regulation and amino acid homeostasis. Human ACE2 acts as the receptor for severe acute respiratory syndrome coronaviruses SARS-CoV and SARS-CoV-2. ACE2 is also widely expressed in epithelial cells of lungs, heart, kidney and pancreas. It is considered an important drug target for treating SARS-CoV-2, as well as pulmonary diseases, heart failure, hypertension, renal diseases and diabetes. Despite the critical importance, the mechanism of ligand binding to the human ACE2 receptor remains unknown. Here, we address this challenge through all-atom simulations using a novel ligand Gaussian accelerated molecular dynamics (LiGaMD) method. Microsecond LiGaMD simulations have successfully captured both binding and unbinding of the MLN-4760 inhibitor in the ACE2 receptor. In the ligand unbound state, the ACE2 receptor samples distinct Open, Partially Open and Closed conformations. Ligand binding biases the receptor conformational ensemble towards the Closed state. The LiGaMD simulations thus suggest a conformational selection mechanism for ligand recognition by the ACE2 receptor. Our simulation findings are expected to facilitate rational drug design of ACE2 against coronaviruses and other related human diseases.

## Introduction

Angiotensin converting enzyme 2 (ACE2) plays a key role in renin-angiotensin system regulation and amino acid homeostasis (Gheblawi et al., 2020;Gross et al., 2020). Human ACE2 acts as the receptor for severe acute respiratory syndrome coronaviruses SARS-CoV and SARS-CoV-2. ACE2 plays a vital role as a catalytic protease converting angiotensin II to angiotensin 1-7 and angiotensin I to angiotensin 1-9. Proteolytic reactions of these peptide hormones aid in the conversion of vasoconstrictors to vasodilators and thus help to maintain blood pressure (Gross et al., 2020). Likewise, ACE2 is widely expressed in epithelial cells of lungs, heart, kidney, and pancreas. It is an important drug target for treating pulmonary diseases, heart failure, hypertension, renal diseases and diabetes (Patel et al., 2016;Yan et al., 2020). ACE2 also stabilizes amino acid transporter B°AT1 to regulate the gut microbiome and amino acid homeostasis (Perlot and Penninger, 2013).

Human ACE2 has been identified as the functional receptor for severe acute respiratory syndrome coronaviruses including SARS-CoV and SARS-CoV-2 (Hoffmann et al., 2020). SARS-CoV-2 is responsible for 2019 coronavirus pandemic (COVID-19). By mid-October 2020, ∼40 million people have been infected by COVID-19 with ∼1.1 million deaths around the world. With the unprecedented pandemic, it is of paramount importance to investigate virus infection and develop effective treatments of SARS-CoV-2. The entry of SARS-CoV-2 is mediated by interaction of the receptor binding domain (RBD) in the virus spike protein S1 subunit with the host ACE2 receptor. The transmembrane protease serine 2 (TMPRSS2) promotes priming of the spike protein and facilitates its S2 subunit to initiate the viral-cell membrane fusion. Hence, inhibiting the interaction between the viral RBD and host ACE2 presents a promising strategy for blocking SAR-COV-2 entry to the human cells.

ACE2 consists of the N-terminal catalytic peptidase domain (PD) on the extracellular side and the C-terminal transmembrane collectrin-like domain (CLD) with a cytoplasmic tail on the intracellular side. The enzyme PD domain can be inhibited by compounds like MLN-4760 (Towler et al., 2004), which bind to the protein active site and prevent substrate binding. MLN-4760 binding biases the receptor to adopt a “Closed” conformation, in which the protein active site formed by two subdomains of the PD is closed from the external environment (Figure 1A). Furthermore, the receptor undergoes conformational changes with hinge-bending movement of the dynamic N-terminal subdomain I relative to the stable subdomain II, e.g., ∼16° bending upon inhibitor binding (Towler et al., 2004). In the absence of ligand binding, the two subdomains move apart from each other and the protein active site becomes exposed to solvent in an “Open” conformation (Figure 1A). In complex with RBD of the SARS-CoV or SARS-CoV-2, the ACE2 receptor also adopts a “Partially Open” conformation, in which the subdomain I lies between the “Open” and “Closed” conformations (Gross et al., 2020) (Figure 1A). Among over 20 experimental structures of the ACE2 receptor present in Protein Data Bank (PDB), most of them exhibit “Open” and “Partially Open” conformations but only one structure has been identified in the “Closed” conformation (PDB: 1R4L) (Towler et al., 2004). Despite tremendous efforts to determine these experimental structures (Huentelman et al., 2004;Towler et al., 2004;Li et al., 2005;Hernández Prada et al., 2008;Song et al., 2018;Shang et al., 2020;Wang et al., 2020;Yan et al., 2020), the dynamics and functional mechanism of the ACE receptor are still poorly understood (Nami et al., 2020).

**Figure 1.**
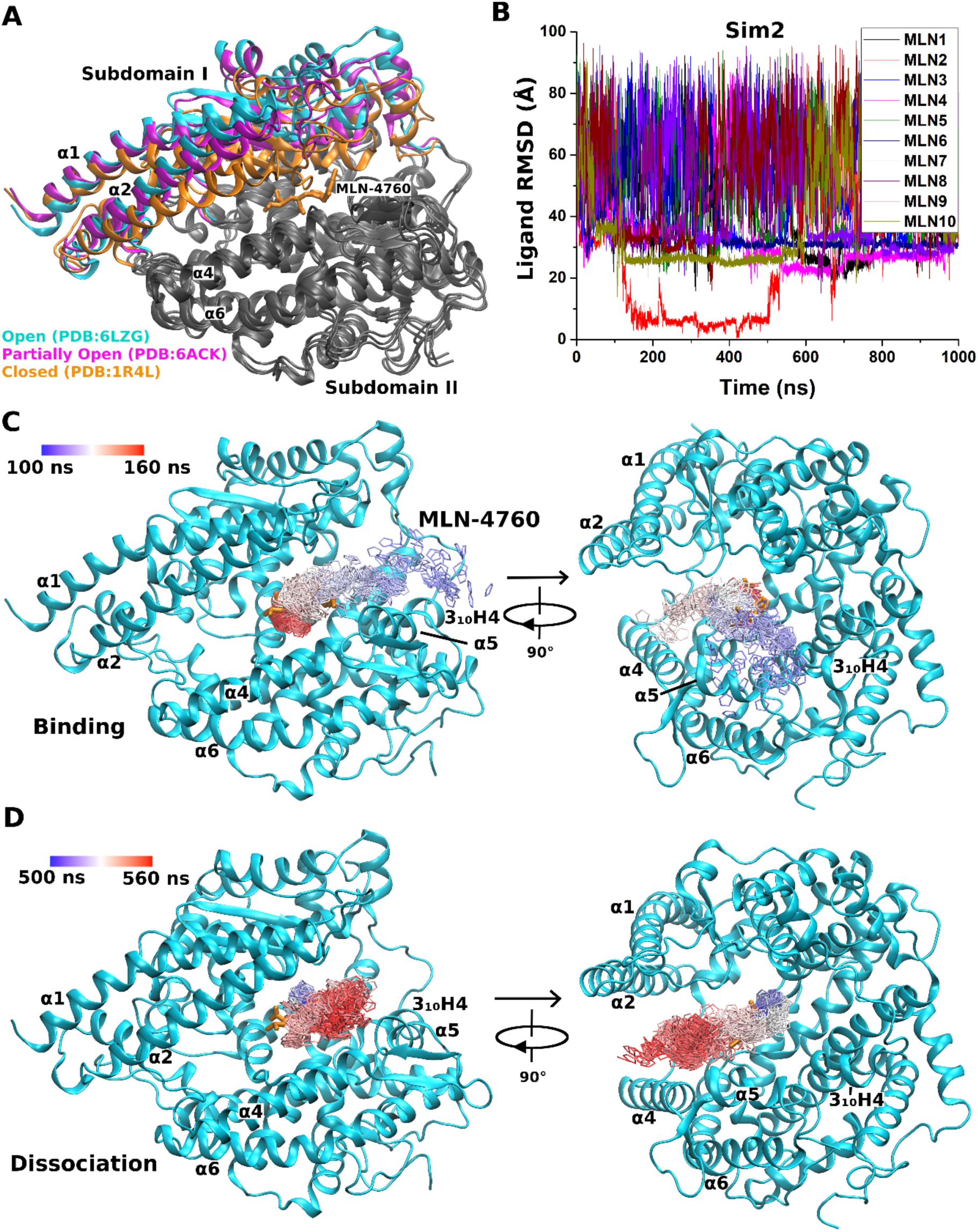
(A) X-ray and cryo-EM strucutres of the ACE2 receptor with subdomain I in the “Open” (cyan, PDB: 6LZG), “Partially Open” (magenta, PDB: 6ACK) and “Closed” (orange, PDB: 1R4L) conformations. Subdomain II is stable and colored in gray. In the “Closed” conformation, the receptor is bound by the MLN-4760 inhibitor. The protein is shown as ribbons and the ligand as sticks (orange). (B) Root-mean-square deviations (RMSDs) of ten MLN-4760 inhibitor molecules relative to the bound X-ray conformation (PDB: 1R4L) are calculated from the 1000 ns “Sim 2” LiGaMD trajectory, in which the ligand bound to the ACE2 active site during ∼100-400 ns and then dissociated into the bulk solvent. (C) Two views of the observed ligand binding pathway, for which the center ring of MLN-4760 is represented by lines and colored by simulation time in a blue-white-red (BWR) color scale. (D) Two views of the observed ligand dissociation pathway, for which the center ring of MLN-4760 is represented by lines and colored by simulation time in a blue-white-red (BWR) color scale.

MLN-4760 is a highly selective and potent (IC_50_: 0.44 nM) small-molecule inhibitor of the ACE2 receptor (Dales et al., 2002). The inhibitor has two carboxylate groups contributing to -2 net charge of the molecule (**Figure S1**). One of the negatively charged carboxylate groups interacts with the positively charged zinc ion, by which the ACE2 receptor functions as a metallopeptidase enzyme. Depending on ligand or viral RBD binding, the receptor adopts different conformations, but the pathways and mechanism of ligand binding in the ACE2 receptor remain unknown. In the context of SARS-CoV-2 and many other medical implications, it is important to understand the mechanism of ligand recognition by the ACE2 receptor in order to design effective drugs against the virus.

Ligand Gaussian accelerated molecular dynamics (LiGaMD) (Miao et al., 2020) is an enhanced sampling computational technique that is developed to efficiently simulate both binding and unbinding of ligand molecules. It is based on Gaussian accelerated molecular dynamics (GaMD), which works by adding a harmonic boost potential to smooth the biomolecular potential energy surface (Miao et al., 2015). GaMD greatly reduces energy barriers and accelerates biomolecular simulations by orders of magnitude (Miao, 2018). GaMD provides unconstrained enhanced sampling without the requirement of pre-defined collective variables or reaction coordinates. Moreover, because the boost potential exhibits a Gaussian distribution, biomolecular free energy profiles can be properly recovered through cumulant expansion to the second order (Miao et al., 2015). In LiGaMD (Miao et al., 2020), the ligand non-bonded interaction potential energy is selectively boosted to enable ligand dissociation. Another boost potential is applied to the remaining potential energy of the entire system in a dual-boost algorithm to facilitate ligand rebinding. LiGaMD has been demonstrated on host-guest and protein-ligand binding model systems. LiGaMD allows us to capture repetitive ligand binding and unbinding, and thus characterize both ligand thermodynamics and kinetics simultaneously. The calculated ligand binding free energy and kinetic rate constants compared very well with experimental data (Miao et al., 2020).

Here, we have applied all-atom LiGaMD simulations to investigate binding and unbinding of the MLN-4760 inhibitor and associated conformational changes of the ACE2 receptor. Microsecond LiGaMD simulations were able to capture both ligand binding and unbinding. The ACE2 receptor sampled Open, Partially Open and Closed conformational states, being consistent with previous experimental structures. Ligand binding could bias the receptor conformational ensemble to the Closed state, suggesting a conformational selection mechanism rather than induced fit. In summary, the LiGaMD simulations allowed us to understand the mechanism of ligand recognition by the ACE2 receptor.

## Materials and Methods

### Ligand Gaussian accelerated molecular dynamics (LiGaMD)

LiGaMD is an enhanced sampling computational technique that is developed to efficiently simulate ligand binding and unbinding based on the previous GaMD (Miao et al., 2015). Details of the LiGaMD method has been described in the previous study (Miao et al., 2020). A brief summary will be provided here.

We consider a system of ligand *L* binding to a protein *P* in a biological environment *E*. The system comprises of *N* atoms with their coordinates 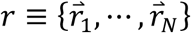 and momenta 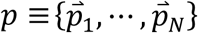. The system Hamiltonian can be expressed as:

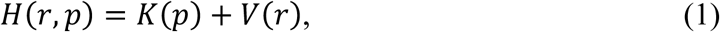

where *K*(*p*) and *V*(*r*) are the system kinetic and total potential energies, respectively. Next, we decompose the potential energy into the following terms:

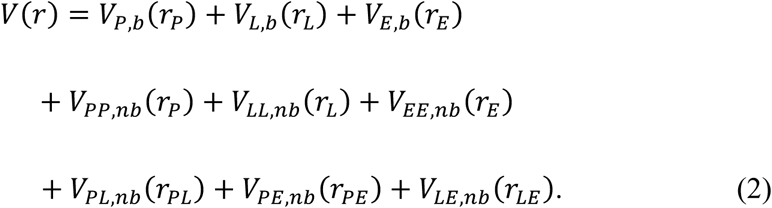

where *V*_*P,b*_, *V*_*L,b*_ and *V*_*E,b*_ are the bonded potential energies in protein *P*, ligand *L* and environment *E*, respectively. *V*_*PP,nb*_, *V*_*LL,nb*_ and *V*_*EE,nb*_ are the self non-bonded potential energies in protein *P*, ligand *L* and environment *E*, respectively. *V*_*PL,nb*_, *V*_*PE,nb*_ and *V*_*LE,nb*_ are the non-bonded interaction energies between *P-L, P-E* and *L-E*, respectively. According to classical molecular mechanics force fields (Cornell et al., 1996;Duan et al., 2003;Vanommeslaeghe et al., 2010;Vanommeslaeghe and MacKerell, 2014), the non-bonded potential energies are usually calculated as:

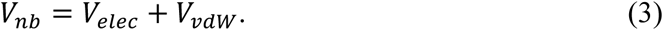

Where *V*_*elec*_ and *V*_*vdw*_ are the system electrostatic and van der Waals potential energies. Presumably, ligand binding mainly involves the non-bonded interaction energies of the ligand, *V*_*L,nb*_(*r*) = *V*_*LL,nb*_(*r*_*L*_) + *V*_*PL,nb*_(*r*_*PL*_) + *V*_*LE,nb*_(*r*_*LE*_). In LiGaMD, we add boost potential selectively to the essential ligand non-bonded potential energy according to the GaMD algorithm:

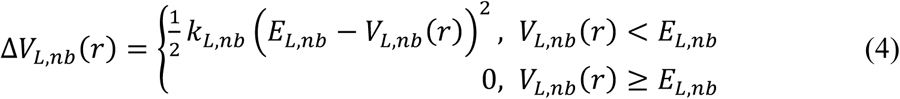

where *E*_*L,nb*_ is the threshold energy for applying boost potential and *k*_*L,nb*_ is the harmonic constant. For simplicity, the subscript of Δ*V*_*L,nb*_(*r*), *E*_*L,nb*_ and *k*_*L,nb*_ is dropped in the following. When *E* is set to the lower bound *E=V*_*max*_, *k*_0_ can be calculated as:

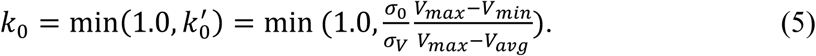

Alternatively, when the threshold energy *E* is set to its upper bound 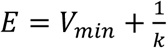, *k*_0_ is set to:

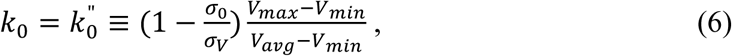

if 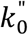 is found to be between *0* and *1*. Otherwise, *k*_0_ is calculated using Eqn. (5).

Next, one can add multiple ligand molecules in the solvent to facilitate ligand binding to proteins in MD simulations (Dror et al., 2011;Miao et al., 2018). This is based on the fact that the ligand binding rate constant *k*_on_ is inversely proportional to the ligand concentration. The higher the ligand concentration, the faster the ligand binds, provided that the ligand concentration is still within its solubility limit. In addition to selectively boosting the bound ligand, another boost potential could thus be applied on the unbound ligand molecules, protein and solvent to facilitate both ligand dissociation and rebinding. The second boost potential is calculated using the total system potential energy other than the non-bonded potential energy of the bound ligand in Eqn. (2) as:

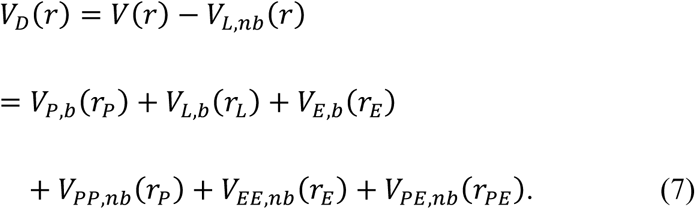

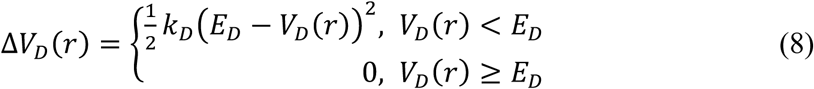

where *E*_*D*_ and *k*_*D*_ are the corresponding threshold energy for applying the second boost potential and the harmonic constant, respectively. This leads to dual-boost LiGaMD (LiGaMD_Dual) with the total boost potential Δ*V*(*r*) = Δ*V*_*L,nb*_(*r*) + Δ*V_D_*(*r*).

### Energetic Reweighting

To calculate potential of mean force (PMF) (Roux, 1995) from LiGaMD simulations, the probability distribution along a reaction coordinate is written as *p*^*^(*A*). Given the boost potential 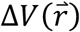 of each frame, *p*^*^(*A*) can be reweighted to recover the canonical ensemble distribution, *p*(*A*), as:

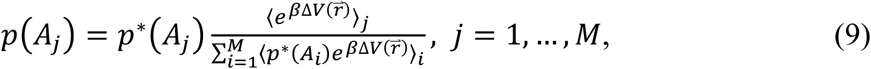

where *M* is the number of bins, *β* = *k*_*B*_*T* and 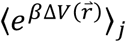 is the ensemble-averaged Boltzmann factor of 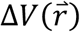 for simulation frames found in the *j*^th^ bin. The ensemble-averaged reweighting factor can be approximated using cumulant expansion:

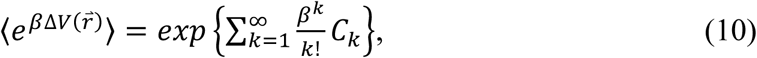

where the first two cumulants are given by

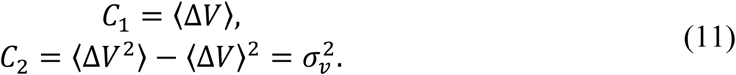

The boost potential obtained from LiGaMD simulations usually follows near-Gaussian distribution. Cumulant expansion to the second order thus provides a good approximation for computing the reweighting factor (Miao et al., 2014;Miao et al., 2015). The reweighted free energy *F*(*A*) = −*k*_B_*T* ln *p*(*A*) is calculated as:

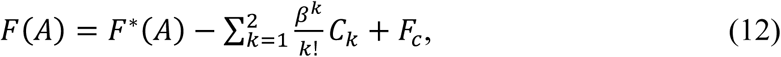

where *F*^*^(*A*) = −*k*_B_*T* ln *p*^*^(*A*) is the modified free energy obtained from LiGaMD simulation and *F_c_* is a constant.

### Simulations of ligand binding in human ACE2 receptor

LiGaMD simulations using the dual-boost scheme were performed on ligand binding to the ACE2 receptor. The X-ray crystal structure of the MLN-4760 inhibitor-bound ACE2 (PDB: 1R4L (Towler et al., 2004)) was used to prepare the simulation system **(Figure S1)**. The N- and C-termini of the peptides were capped with the acetyl (ACE) and N-methyl amide (NME) neutral groups, respectively. The missing hydrogen atoms were added using the *tleap* module in AMBER (Case et al., 2005a). The AMBER ff19SB force field (Tian et al., 2019) were used for the protein. The MLN-4760 ligand was modeled with GAFF-2 (Case et al., 2020) force field. Atomic partial charges of ligand were obtained through B3LYP/6-31G* quantum calculations of the electrostatic potential, for which the charges were fitted using the antechamber program (Case et al., 2005a;Case et al., 2005b). Each system was neutralized by adding counter ions and immersed in a cubic TIP3P (Jorgensen et al., 1983) water box, which was extended 10 Å from the receptor surface. A total of 10 ligand molecules (one in the X-ray bound conformation and another nine placed randomly in the solvent) were included in the system to facilitate ligand binding. This design was based on the fact that the ligand binding rate is inversely proportional to the ligand concentration. The higher the ligand concentration, the faster the ligand binds, provided that the ligand concentration is still within its solubility limit.

The built simulation system was energy minimized with 1 kcal/mol/Å^2^ constraints on the heavy atoms of the protein and ligand, including the steepest descent minimization of 5,000 steps followed by a conjugate gradient minimization of 5,000 steps. The system was then heated from 0 K to 300 K for 200 ps. It was further equilibrated using the NVT ensemble at 300K for 800 ps and the NPT ensemble at 300 K and 1 bar for 1 ns with 1 kcal/mol/Å^2^ constraints on the heavy atoms of the protein and ligand, followed by 2 ns short cMD without any constraint. The LiGaMD simulations proceeded with 14 ns short cMD to collect the potential statistics, 64 ns GaMD equilibration after adding the boost potential and then three independent 1000 ns production runs **(Table S1)**. Initial testing simulations showed that when the threshold energy for applying boost potential to the ligand non-bonded energy was set to the lower bound (i.e., *E* = *V*_max_), the bound ligand maintained the X-ray conformation and did not dissociate. In comparison, when the threshold energy was set to the upper bound (i.e., *E* = *V*_min_+1/*k*), it enabled high enough boost potential to dissociate the ligand from the protein. In addition, boost applied not only to the ligand non-bonded energy but also to all the bonded energies can accelerate the molecular transitions for ligand dissociation and protein conformational change. Therefore, the threshold energy for applying the ligand boost potential was set to the upper bound in the LiGaMD_Dual simulations. Similarly, second boost potential was applied to the system total potential energy other than the ligand bonded/non-bonded potential energy to provide sufficient acceleration for sampling ligand rebinding. The threshold energy was set to the upper bound for rebinding part as well. LiGaMD_Dual production simulation frames were saved every 0.2 ps for analysis.

The VMD (Humphrey et al., 1996) and CPPTRAJ (Roe and Cheatham, 2013) tools were used for simulation analysis. The 2D PMF profiles were calculated through energetic reweighting of the LiGaMD_Dual simulations using *PyReweighting* toolkit (Miao et al., 2014). 2D PMF profiles of the interdomain Glu56:CA - Ser128:CA atom distance and root-mean-square deviation (RMSD) of the MLN-4760 inhibitor relative to X-ray conformation were calculated to analyze conformational changes of the protein upon ligand binding. Residues Glu56 and Ser128 lie on the tip of α4 (subdomain I) and α6 helices (subdomain II), respectively. The bin size was set to 2.0 Å for the atom distances. Similarly, 2D PMF profiles of the Glu56:CA - Ser128:CA atom distance and RMSD of subdomain I relative to X-ray conformation were calculated to characterize domain flexibility in the ACE2 receptor. The cutoff of simulation frames in one bin for 2D PMF reweighting was set to 500. Structural clustering was performed on the LiGaMD simulations based on the RMSD of the receptor as well as the ligand snapshots using hierarchical agglomerative algorithm in CPPTRAJ (Roe and Cheatham, 2013). The RMSD cutoff was set to 2.0 A. The top structural clusters were identified as the representative conformations of the receptor and ligand corresponding to low-energy states in the PMF profiles.

## Results

Atomic trajectores were obtained from LiGaMD simulations on the MLN-4760 inhibitor-bound ACE2 receptor (PDB: 1R4L) (Towler et al., 2004) (Figure 1). During the LiGaMD equlibration, the bound ligand dissociated from the active site to the bulk solvent, accompanied by large conformational changes of the protein subdomain I **(Figure S2)**. Upon ligand dissociation, subdomain I of the receptor changed from the “Closed” to “Open” conformation. Three independent 1000 ns LiGaMD production runs were further performed with randomized intitial atomic velocities after the equilbration simulation.

During one of the three 1000 ns LiGaMD production simulations (“Sim 2” in **Table S1**), the MLN-4760 inhibitor bound to the active site of the ACE2 receptor during ∼100-500 ns (Figure 1B), while no ligand binding was observed in the other two simulations **(Figure S3)**. In “Sim 2”, RMSD of the ligand relative to the 1R4L X-ray structure reached a minimum of ∼0.99 Å near ∼420 ns (Figure 1B). The ligand then dissociated at ∼500 ns into the bulk solvent and stayed unbound during the remainder of the 1000 ns LiGaMD simulation. Meanwhile, the receptor underwent large-scale conformational changes and sampled different conformations inlcuding the “Open”, “Partially Open” and “Closed” (Figure 1A), which were consistent with the receptor experimental structures.

### LiGaMD simulations captured both ligand binding and dissociation of ACE2

During the “Sim 2” LiGaMD trajectory, starting from the bulk solvent, one of the MLN-4760 inhibitor molecules first attached to the interface between the receptor 3_10_ H4 and α5 helices within ∼100 ns, moved up into the space between the two protein subdomains and entered the active site of the ACE2 receptor between ∼100-160 ns (Figure 1C). The ligand bound at the active site of the receptor during ∼100-500 ns. At ∼500 ns, the ligand dissociated from the active site to bulk solvent (Figure 1D). The dissociation pathway was observed to be different from that of binding. Ligand dissociated from the opening between the receptor α2 and α4 helices as the subdomain I transitioned from “Closed” to “Open” conformation (Figure 1D). On the other hand, the ligand bound through the space just above the 3_10_ H4 and α5 helices (Figure 1C). During ligand binding and dissociation in the “Sim 2” LiGaMD trajectory, subdomain I of the receptor sampled different conformations. However, such conformational changes were also observed in other two simulations (“Sim1” and “Sim 2”) regardless of ligand binding/dissociation **(Figure S4)**.

### Free energy profiles of ligand binding in the human ACE2 receptor

A 2D potential of mean force (PMF) free energy profile was calculated with the ligand RMSD relative to X-ray conformation and the interdomain distance by combining the three independent 1000 ns LiGaMD production trajectories (Figures 2A). Six low-energy conformational states of the receptor were identified from the PMF profile, including the “Bound (B)”, “Intermediate-1 (I-1)”, “Intermediate-2 (I-2)”, “Intermediate-3 (I-3)”, “Unbound-1 (U-1)” and “Unbound-2 (U-2)”. Particularly, the system adopted the “Bound” state with ligand RMSD < 5 Å, the “Unbound” state with ligand RMSD > 35 Å and intermediatestates with 5 – 35 Å ligand RMSD relative to the 1R4L X-ray structure.

**Figure 2.**
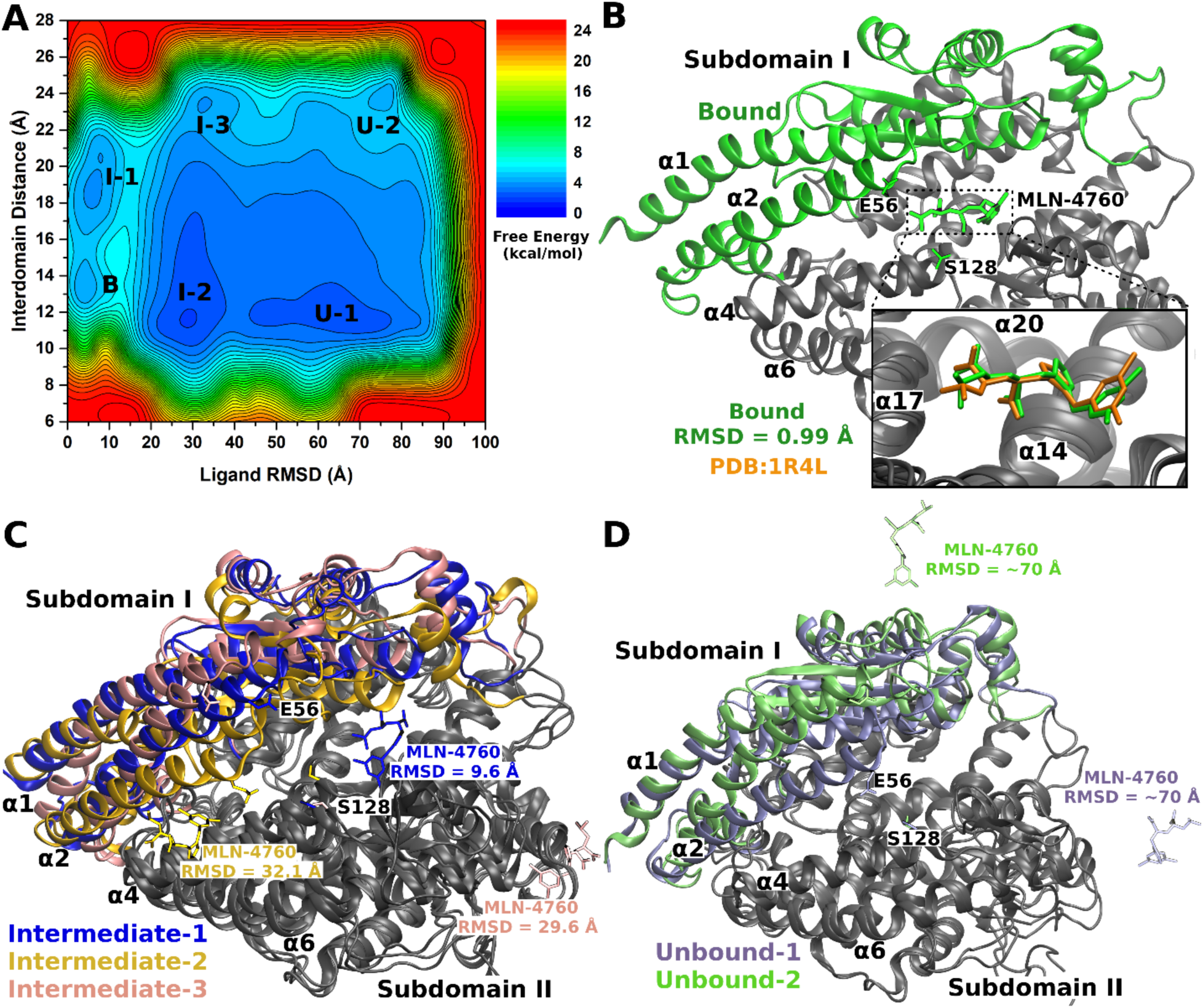
(A) The 2D potential of mean force (PMF) free energy profile of the ligand RMSD and interdomain distance calculated by combining the three independent 1000 ns LiGaMD production simulations of human ACE2 receptor. Six low energy conformational states were identified, including the “Bound (B)”, “Intermediate-1 (I-1)”, “Intermediate-2 (I-2)”, “Intermediate-3 (I-3)”, “Unbound-1 (U-1)” and “Unbound-2 (U-2)”. (B) Conformations of the ACE2 receptor and MLN-4760 ligand in the “Bound (B)” state (green) compared with the X-ray conformation (Orange, PDB: 1R4L). (C) Conformations of the ACE2 receptor during binding of MLN-4760 in the “Intermediate-1 (I-1)” (blue) and “Intermediate-2 (I-2)” (yellow) and “Intermediate-3 (I-3)” (pink) states. (D) Conformations of the ACE2 receptor in the “Unbound-1 (U-1)” (ice blue) and “Unbound-2 (U-2)” (lime) states. The protein is shown as ribbons and ligand as sticks.

In the “Bound” state, the ligand bound at the protein active site and the protein interdomain distance was ∼14 Å. The ligand exhibited a minimum RMSD of 0.99 Å compared with the X-ray structure (Figures 2A **and** 2B). The system sampled three different intermediate states during ligand binding, *i*.*e*., “Intermediate-1 (I-1)”, “Intermediate-2 (I-2)” and “Intermediate-3 (I-3)”. In the I-1 state, the ligand RMSD was ∼9.6 Å and the interdomain distance was ∼18-20 Å (Figures 2A **and** 2C). The ligand was located near the active site making interactions with residues of the α4 helix, α5 helix, α11 helix, 3_10_H8 helix, β4-β5 loop, and α8-3_10_H4 loop in the two protein subdomains. In the “I-2” state, the ligand RMSD was ∼32.1 Å and the interdomain distance was ∼12 Å (Figures 2A **and** 2C). The ligand interacted with the α2 helix of subdomain I and the α4 helix of subdomain II in the receptor. In the “I-3” state, the ligand RMSD was ∼29.6 Å and the interdomain distance was ∼23 Å (Figures 2A **and** 2C). The ligand interacted with the α8 helix, α20 helix, α8-β3 loop, and the C-terminal loop after the α20 helix in the protein subdomain II (Figure 2C). The system sampled two “Unbound (U-1)” and “Unbound (U-2)” states, where the ligand RMSD was ∼60-70 Å and the interdomain distances were ∼12 Å and ∼23 Å, respectively. In these states, the ligand was found far away from the receptor in the bulk solvent and the receptor could change between the “Closed” and “Open” conformations (Figures 2A **and** 2D).

### Low-energy intermediate conformational states of ligand binding to human ACE2 receptor

Low-energy intermediate conformational states “I-1”, “I-2” and “I-3” of the MLN-4760 inhibitor binding to the human ACE2 receptor identified from the LiGaMD simulation free energy profiles are shown in Figures 3A, 3B **and** 3C, respectively. Polar and charged groups present in different parts of the receptor made favorable interactions with the charged carboxylate groups and the polar chloride and nitrogen atoms in the ligand molecule. These interactions played important role in recognition and binding of the MLN-4760 inhibitor to the receptor.

**Figure 3.**
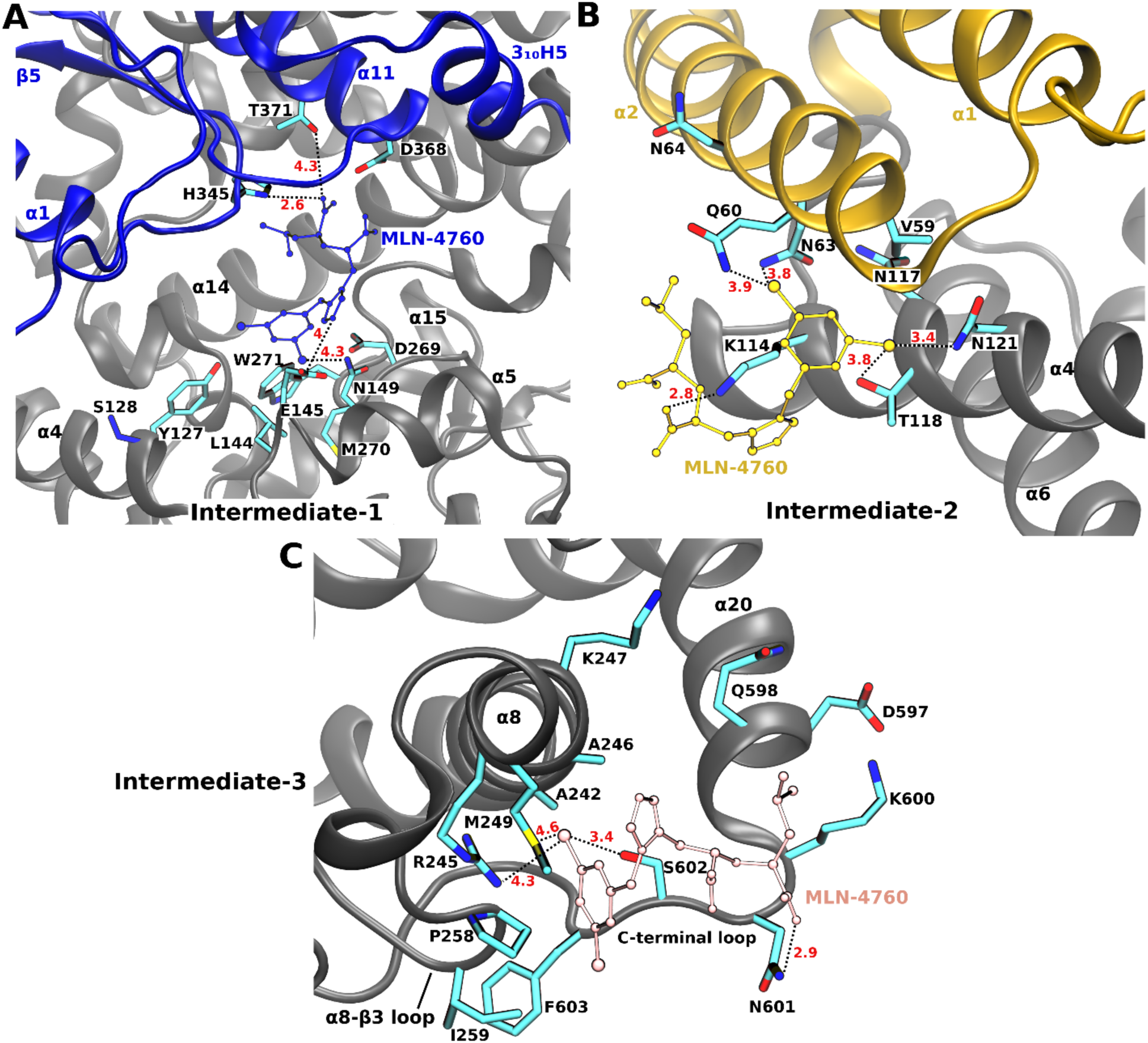
(A) The “Intermediate-1 (I-1)” conformational state of the MLN-4760 ligand (blue balls and sticks) bound to the ACE2 receptor (ribbons). Residues of subdomain I (blue) and subdomain II (gray) including H345, T371, E145 and N149 formed polar interactions with the ligand. (B) The “Intermediate-2 (I-2)” conformational state of MLN-4760 (yellow) bound to the ACE2 receptor (ribbons). Residues of subdomain I (yellow) and subdomain II (gray) including Q60, N63, N171 and N121 formed polar interactions with the ligand. (C) The “Intermediate-3 (I-3)” conformational state of MLN-4760 (pink) bound to the ACE2 receptor (ribbons). Residues of subdomain I (pink) and subdomain II (gray) including R245, M249, S602 and N601 formed polar interactions with the ligand.

In the intermediate “I-1” state, the receptor adopted a “Partially Open” conformation with ∼18 Å interdomain distance. The ligand molecule was located near the recetpor active site with ∼9.6 Å RMSD relative to the X-ray structure and formed interactions with residues from both subdomains of ACE2. One of the ligand carboxylate groups formed ionic interaction with the positively charged protein residue His345 (Figure 3A). The carboxylate group also interacted with protein residue Thr371. Similary, one of the nitrogen atoms in the ligand’s central ring formed ionic interaction with the negatively charged protein residue Glu145. The ligand chloride group formed polar interactions with protein reisude Asn149.

In the intermediate “I-2” state, the receptor adopted the “Closed” conformation and the ligand interacted with the α2 helix of subdomain I and the α4 helix of subdomain II in the receptor (Figure 3B). One of the ligand chloride atoms formed polar interactions with protein residues Gln60 and Asn63, while the other chloride atom formed polar interactions with protein residues Thr118 and Asn121. One of the ligand’s negatively charged carboxylate groups formed ionic interaction with the protein postively charged residue Lys114. The ligand terminal benzene ring formed hydrohpobic interactions with protein residue Val59 (Figure 3B).

In the intermediate “I-3” state, one of the ligand chloride atoms formed polar interactions with protein residues Arg245, Ser602 and Met249 of subdomain II (Figure 3C). One of the ligand charged caroxylate groups formed polar interactions with protein residue Asn601. The ligand terminal benzene ring formed hydrohpobic interactions with Pro258 in the receptor.

During one of the LiGaMD trajectories (“Sim3”), the MLN-4760 ligand was observed to interact with residue Gly354 of the ACE2 receptor **(Figure S5)**. The ligand terminal methyl group was able to form close hydrophobic interactions with the Gly354 residue in subdomain I of the ACE2 receptor. Notably, residue Gly354 is considered an important residue making direct interactions with the RBD of SARS-CoV and SARS-CoV-2 as seen in the experimental structures (Wang et al., 2020).

### Conformational plasticity of human ACE2 receptor

In the LiGaMD simulations, while the protein subdomain II was stable maintaining the 1R4L X-ray conformation with ∼2-4 Å RMSD **(Figures S2 and 4B)**, subdomain I in the human ACE2 receptor exhibited high flexibility and underwent large conformational changes with ∼3-10 Å RMSD compared with the X-ray conformation (Figure 4A). We calculated 2D PMF regarding the interdomain distance and subdomain I RMSD relative to the 1R4L X-ray conformation. Three low-energy conformational states were identified in the PMF profile, including the “Open”, “Partially Open” and “Closed” (Figure 3C).

**Figure 4.**
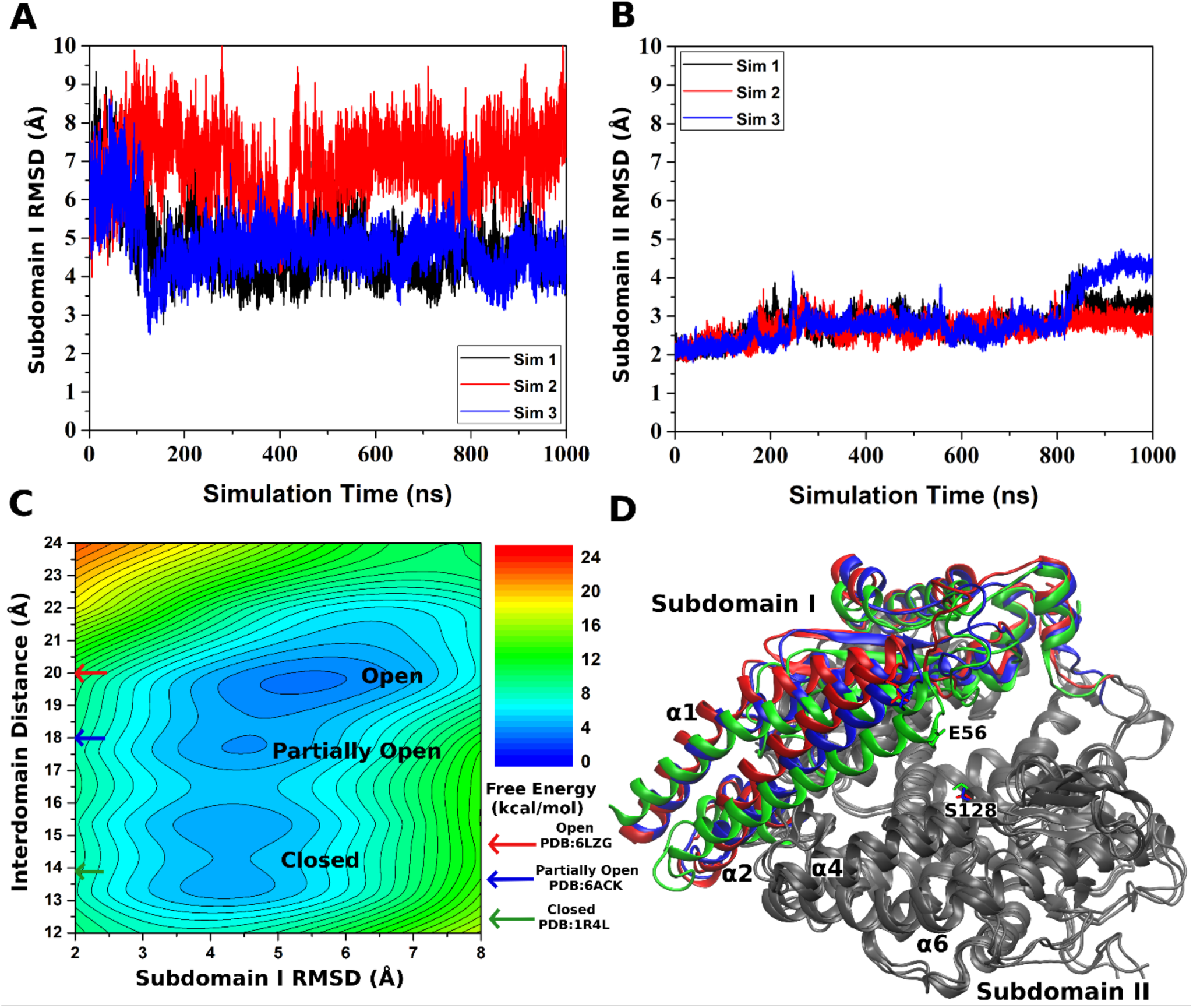
RMSDs of (A) subdomain I and (B) subdomain II of the ACE2 receptor relative to the closed X-ray conformation (PDB:1R4L) are calculated from three independent LiGaMD production simulations. (C) 2D potential of mean force (PMF) of the subdomain I RMSD and interdomain distance calculated by combining the three LiGaMD simulations. Three low energy conformational states of the receptor are identified in the PMF profile, including the “Closed”, “Partially Open” and “Open”, which are similar to the 6LZG, 6ACK and 1R4L PDB structures, respectively. (D) Low-energy conformations of the ACE2 receptor with subdomain I found in the “Open” (red), “Partially Open” (blue) and “Closed” (green) states in the LiGaMD simulations. Subdomain II is stable and colored in gray.

In the “Closed” conformation, subdomain I moved near subdomain II closing the acitve site. The receptor interdomain distance was ∼14 Å and RMSD of subdomain I was ∼4 Å compared with the 1R4L X-ray structure (Figure 3C). In the “Partially Open” conformation, the receptor interdomain distance increased to ∼18 Å and RMSD of subdomain I was ∼4 Å compared with the 1R4L X-ray structure (Figure 3C). Finally, the receptor interdomain distance could increase further to ∼20 Å and the subdomain I RMSD relative to the 1R4L X-ray structure increased to ∼6 Å in the “Open” conformation. Notably, conformations of the ACE2 receptor in the “Partially Open” and “Open” low-energy states were closely similar to the experimental 6ACK cryo-EM and 6LZG X-ray structures, respectively (Figure 3C). Therefore, the different low-energy states of ACE2 receptor revealed from our LiGaMD simulations highlighted the receptor conformational plasticity during its function for ligand binding and interactions with other proteins (e.g., the coronavirus spike protein).

## Discussion

Since its discovery in 2000 (Donoghue et al., 2000), the ACE2 receptor has been recognized as a critical protease enzyme with multiple physiological roles in renin-angiotensin system, amino acid transport, gut microbiome ecology and innate immunity. The ACE2 receptor has also been identified as the functional receptor for SARS-CoV and SARS-CoV-2. The COVID-19 pandemic caused by SARS-CoV-2 has been recognized as a serious global health threat as it has no proper treatment and continues to spread across the world. With the infection cases rising daily, it is critical to develop therapeutics against SARS-CoV-2. Here, we have applied all-atom simulations using a novel LiGaMD method to investigate the mechanism of ligand binding to the human ACE2 receptor.

Through LiGaMD enhanced sampling simulations, we have, for the first time, successfully captured both binding and dissociation of a ligand in the human ACE2 receptor. During the simulations, the receptor could sample distinct conformational states, revealing remarkable conformational plasticity of the receptor. Ligand binding biased the receptor conformational ensemble to the Closed state, suggesting a conformational selection mechanism rather than induced fit. Furthermore, the MLN-4760 ligand molecule with -2 net charge, formed polar interactions with charged and polar residues in the low energy intermediate states. This finding suggested that electrostatic interactions played an important role in the recognition and binding/dissociation of the MLN-4760 inhibitor to the ACE2 receptor, being consistent with previous findings of “electrostatic steering” in recognition of charged ligands by proteins (Wade et al., 1998;Miao et al., 2020).

Despite our encouraging simulation findings, it is important to note that the number of ligand binding and unbinding events captured in the presented LiGaMD simulations was too small and the simulation free energy profiles were not converged. More sufficient sampling would be needed in order to obtain converged simulations and calculate the ligand binding thermodynamics and kinetics. This can be potentially achieved through additional and longer simulations, as well as further method developments combining LiGaMD with other enhanced sampling algorithms such as replica exchange (Huang et al., 2018;Oshima et al., 2019) and Markov state models (Plattner and Noe, 2015).

In this context, the MLN-4760 inhibitor binds to the human ACE2 receptor with high affinity (IC_50_: 0.44 nM) (Towler et al., 2004). It is extremely difficult to simulate the ligand dissociation and binding with long-timescale cMD and even the enhanced sampling methods. Very few computational studies have been carried out on ligand binding to the ACE2 receptor. A recent study (Nami et al., 2020) showed that the MLN-4760 ligand could dissociate from the ACE2 receptor upon binding of the viral RBD. However, this study was not able to characterize ligand binding to the ACE2 receptor. In comparison, our LiGaMD simulations could capture both ligand binding and dissociation in the human ACE2 receptor.

In case of SARS-CoV-2 infection, RBD of the viral spike protein attaches to the top region of subdomain I in the ACE2 receptor, which is distant from the receptor peptidase active site. Thus, the MLN-4760 inhibitor does not compete directly with the RBD of SARS-CoV-2. However, there have been evidences of allosteric signaling between the RBD binding site and active site in the ACE2 receptor. Huentelman et. al. (Huentelman et al., 2004) performed *in silico* molecular docking studies to identify a novel ACE2 inhibitor that could also block SARS-CoV spike protein-mediated receptor attachment. Our LiGaMD simulations further highlighted the dynamic nature of the receptor in terms of the large-scale movement of subdomain I and ligand binding. Similarly, LiGaMD simulations showed that MLN-4760 could form hydrophobic interactions with RBD binding site in subdomain I of the receptor. This information could be helpful in designing drugs that could block RBD binding to the receptor. Likewise, since the human ACE2 receptor shows conformational selection for ligand binding, virtual screening using ensemble docking(Lin et al., 2002;Amaro et al., 2008;Amaro et al., 2018) with receptor structural ensembles generated from the LiGaMD simulations will be a promising approach to designing potent drug molecules of the ACE2 receptor.

In summary, we have successfully simulated both ligand binding and dissociation in the human ACE2 receptor using the novel LiGaMD enhanced sampling method. During the LiGaMD simulations, the receptor could sample distinct Closed, Partially Open and Open conformational states. Ligand binding biased the receptor towards the Closed conformation, suggesting a conformational selection mechanism. Our simulation findings are expected to facilitate rational drug design targeting ACE2 for the therapeutic treatments of COVID-19 and other related human diseases.

## Author Contributions

Conceived the study: Y.M., A.B. and S.P.; Performed simulations: A.B.; Analyzed data: A.B. and S.P.; Wrote the manuscript: A.B., S.P. and Y.M.

## Acknowledgements

This work used supercomputing resources with allocation award TG-MCB180049 through the Extreme Science and Engineering Discovery Environment (XSEDE), which is supported by National Science Foundation grant number ACI-1548562, and the Research Computing Cluster at the University of Kansas. This work was supported in part by the National Institutes of Health (R01GM132572) and the startup funding in the College of Liberal Arts and Sciences at the University of Kansas.

## Supplementary Material

The Supplementary Material for this article can be found online at https://www.frontiersin.org/.

