## Supplementary Material for "Mechanism of ligand recognition by human ACE2 receptor"

**Table S1:** Summary of LiGaMD simulations performed on the ACE2 receptor in the presence of the MLN-4760 inhibitor.

| Simulation | $^{a}N_{atoms}$ | Dimension (Å$^3$) | Simulation (ns) | $^{b}\Delta V_{avg}$ (kcal/mol) | $^{c}\sigma_{\Delta v}$ (kcal/mol) |
|---|---|---|---|---|---|
| **Sim1** | 100,449 | 129 x 93 x 93 | 1000 | 67.29 | 9.10 |
| **Sim2** | 100,449 | 129 x 93 x 93 | 1000 | 67.26 | 9.06 |
| **Sim3** | 100,449 | 129 x 93 x 93 | 1000 | 67.27 | 9.10 |

$^{a}N_{atoms}$ is the number of atoms in the simulation system.

$^{b}\Delta V_{avg}$ and $^{c}\sigma_{\Delta V}$ are the average and standard deviation of the LiGaMD boost potential, respectively.

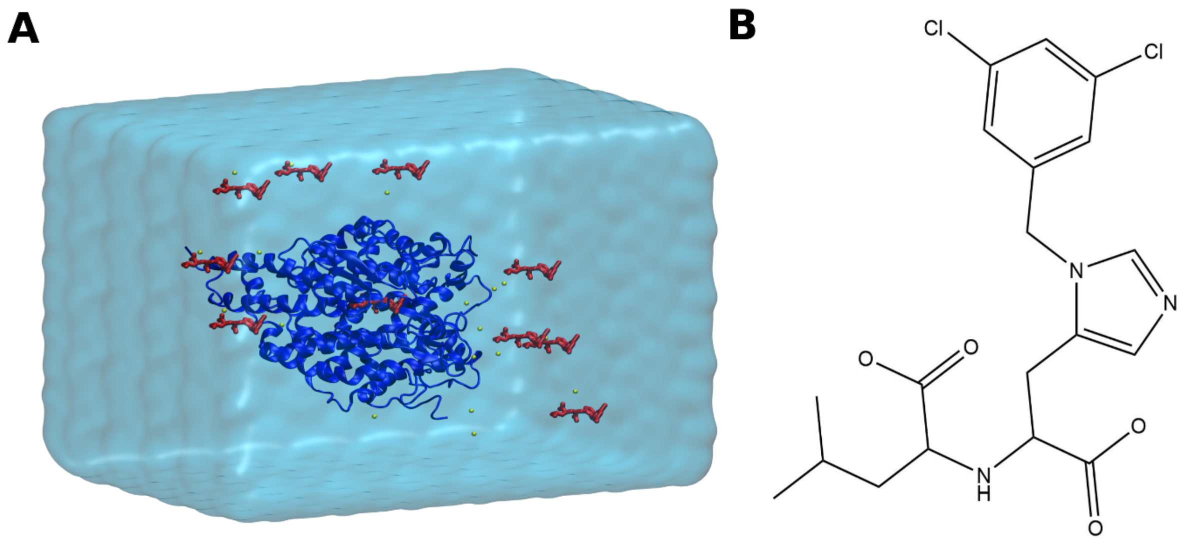


**Figure S1:** (A) Computational model of the ACE2 receptor (blue ribbons) with 10 MLN-4760 ligand molecules (red sticks) (one in the X-ray bound conformation and another nine placed randomly in the solvent) used in LiGaMD simulations. The system was neutralized by adding counter ions and immersed in a cubic TIP3P water box, which was extended 10 Å from the receptor surface. (B) Structure of the MLN-4760 inhibitor molecule.

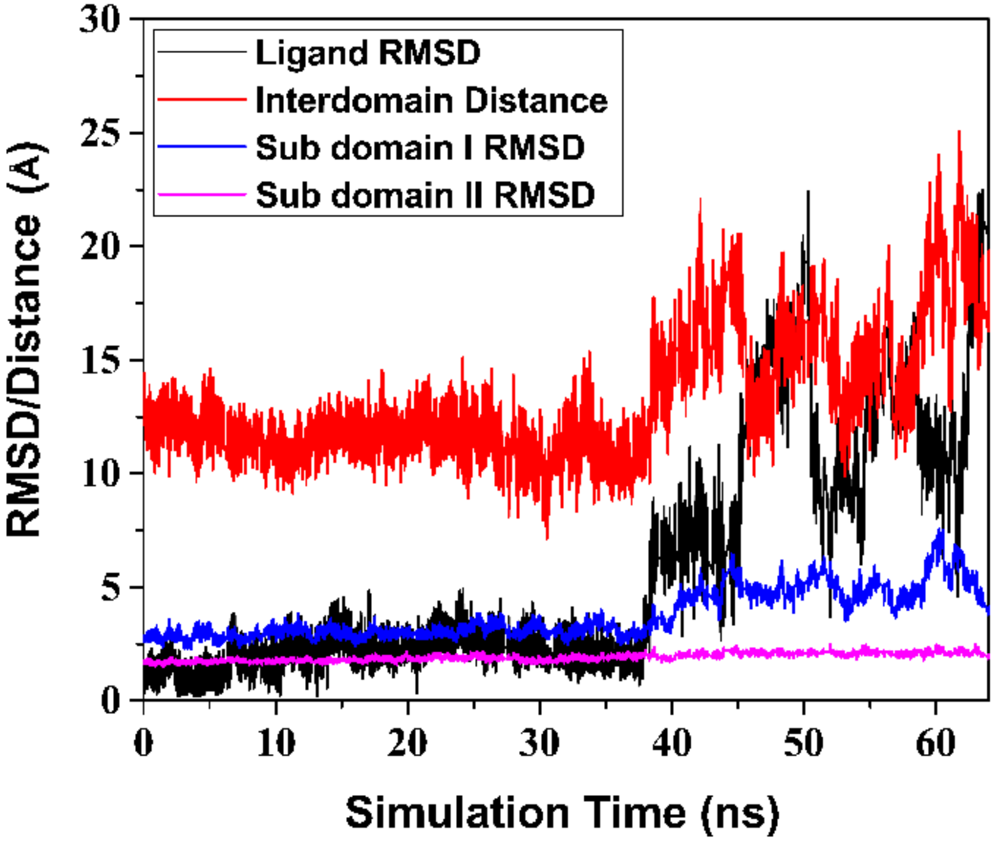


**Figure S2:** Time course of the MLN-4760 ligand RMSD (black), interdomain distance (Glu56:CA – Ser128:CA) (red), sub domain I RMSD (blue) and sub domain II RMSD (pink) calculated from LiGaMD equilibration trajectory, in which the MLN-4760 ligand dissociated from the active site of the receptor.

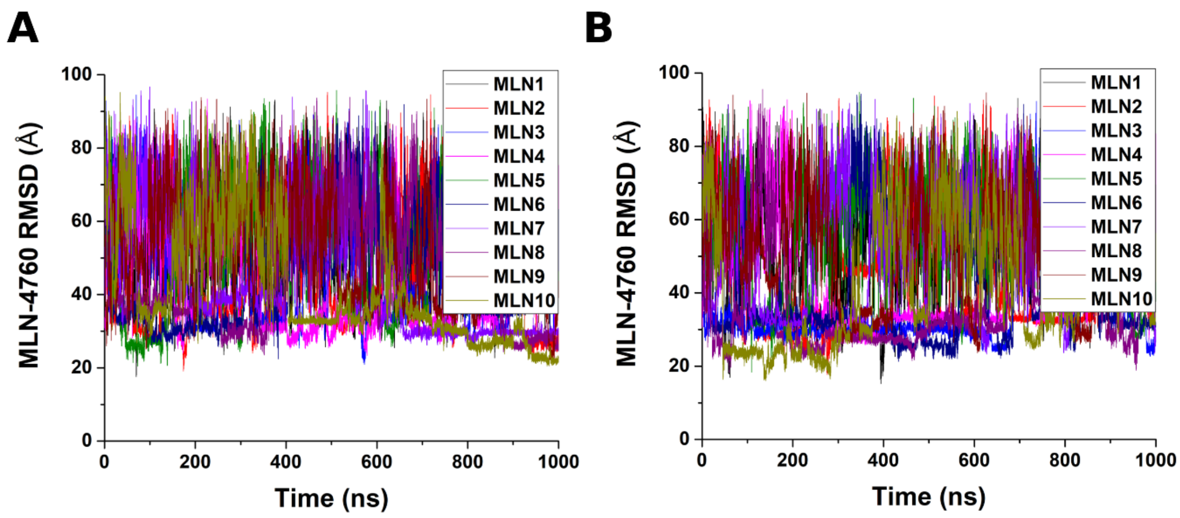


**Figure S3:** Root-mean-square deviations (RMSDs) of the MLN-4760 ligand relative to the bound X-ray conformation (PDB: 1R4L) is plotted for the (A) "Sim 1" and (B) "Sim 3" LiGaMD production trajectories.

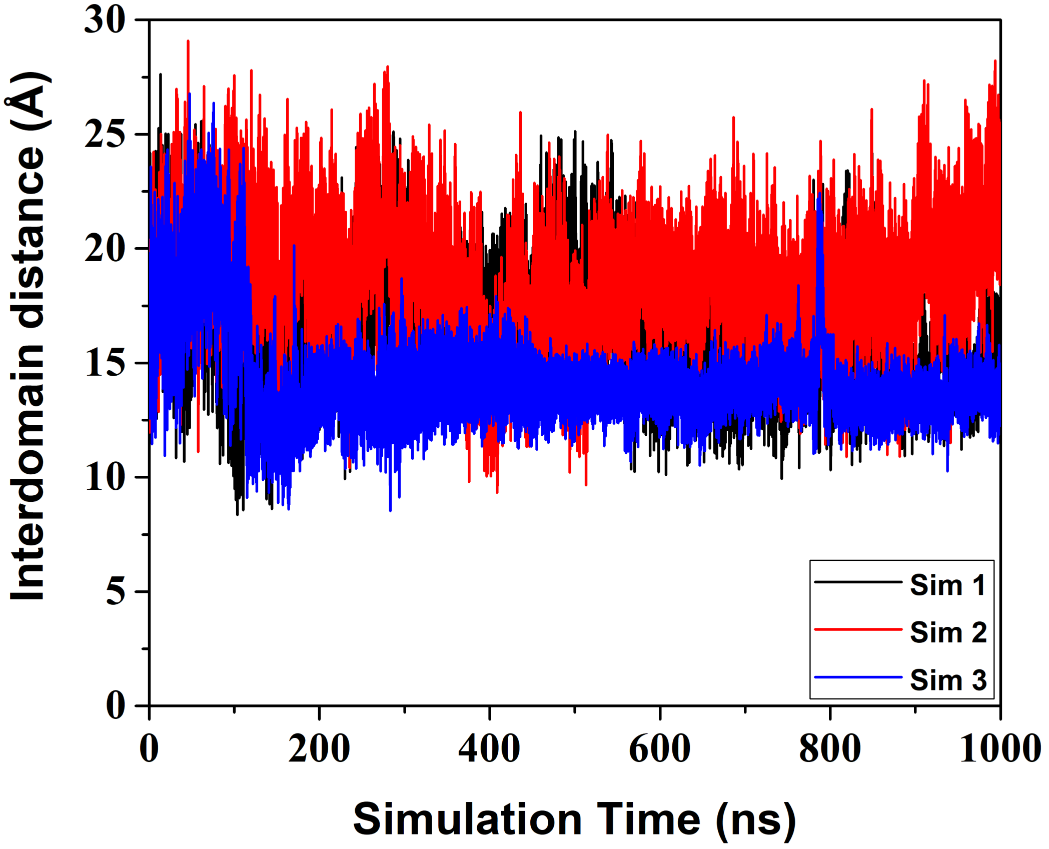


**Figure S4:** Time course of the interdomain distance (Glu56:CA – Ser128:CA) calculated from three independent LiGaMD trajectories including "Sim 1" (black), "Sim 2" (red) and "Sim 3" (blue).

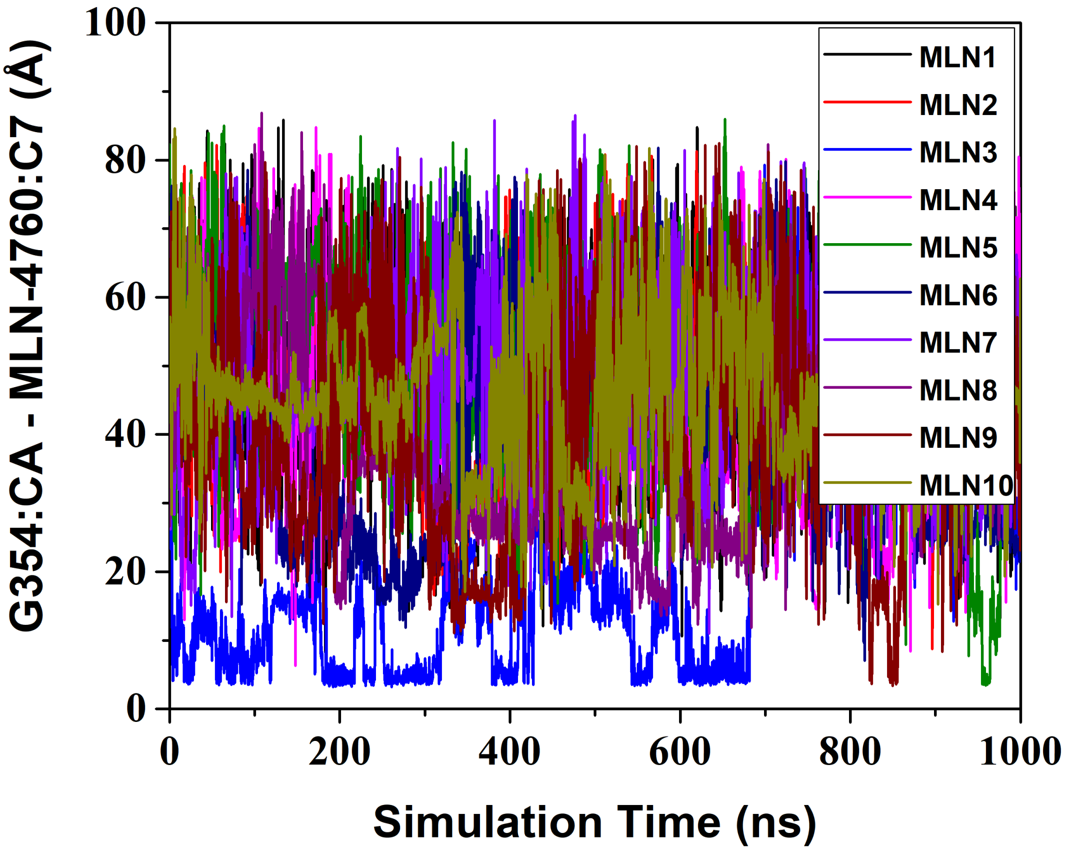


**Figure S5:** Time courses of the Gly354:CA – MLN-4760:C7 distance calculated from "Sim 3" LiGaMD production trajectory. The MLN-4760 ligand formed close hydrophobic interactions with the Gly354 residue in subdomain I of the ACE2 receptor with ~3.5 Å distance between Gly354:CA – MLN-4760:C7 .
